# Developmental, cellular, and biochemical basis of transparency in the glasswing butterfly *Greta oto*

**DOI:** 10.1101/2020.07.02.183590

**Authors:** Aaron F. Pomerantz, Radwanul H. Siddique, Elizabeth I. Cash, Yuriko Kishi, Charline Pinna, Kasia Hammar, Doris Gomez, Marianne Elias, Nipam H. Patel

## Abstract

Numerous species of Lepidoptera have transparent wings, which often possess scales of altered morphology and reduced size, and the presence of membrane surface nanostructures that dramatically reduce reflection. Optical properties and anti-reflective nanostructures have been characterized for several ‘clearwing’ Lepidoptera, but the developmental basis of wing transparency is unknown. We apply confocal and electron microscopy to create a developmental time-series in the glasswing butterfly, *Greta oto*, comparing transparent and non-transparent wing regions. We find that scale precursor cell density is reduced in transparent regions, and cytoskeletal organization differs between flat scales in opaque regions, and thin, bristle-like scales in transparent regions. We also reveal that sub-wavelength nanopillars on the wing membrane are wax-based, derive from wing epithelial cells and their associated microvillar projections, and demonstrate their role in enhancing-anti-reflective properties. These findings provide insight into morphogenesis of naturally organized micro- and nanostructures and may provide bioinspiration for new anti-reflective materials.

## Introduction

The wings of butterflies and moths (Lepidoptera) have inspired studies across a variety of scientific fields, including evolutionary biology, ecology, and biophysics (*1–3*). Lepidopteran wings are generally covered with rows of flat, partially overlapping scales that endow the wings with colorful patterns. Adult scales are chitin-covered projections that serve as the unit of color for the wing. Each scale can generate color through pigmentation via molecules that selectively absorb certain wavelengths of light, structural coloration, which results from light interacting with the physical nanoarchitecture of the scale, or a combination of both pigmentary and structural coloration *(4, 5)*. Cytoskeletal dynamics, including highly organized F-actin filaments during scale cell development, play essential roles in wing scale elongation and prefigure aspects of scale ultrastructure *(6, 7)*.

In contrast to typical colorful wings, numerous species of butterflies and moths possess transparent wings that allow light to pass through, so that objects behind them can be distinctly seen (Fig. 1A-H, *8–10*). This trait has been interpreted as an adaptation in the context of camouflage, in which some lineages evolved transparent wings as crypsis to reduce predation (*11–13*). Transparency results from the transmission of light across the visible spectrum through a material, in this case the chitin membrane, without appreciable absorption or reflection. Levels of reflection are largely determined by the differences in refractive indices between biological tissues and the medium, and a larger difference results in higher surface reflection. Our knowledge on mechanisms underlying transparency in nature is primarily from aquatic organisms, which are frequently transparent, aided by the close match between the refractive indices of their aqueous tissue and the surrounding media — water (*14*). By contrast, transparency is rare and more challenging to achieve on land, primarily due to the large difference between the refractive indices of terrestrial organism’s tissue (*n* = ∼1.3-1.5) and air (*n* = 1), which results in significant surface reflection (*9, 15, 16*).

**Fig. 1.**
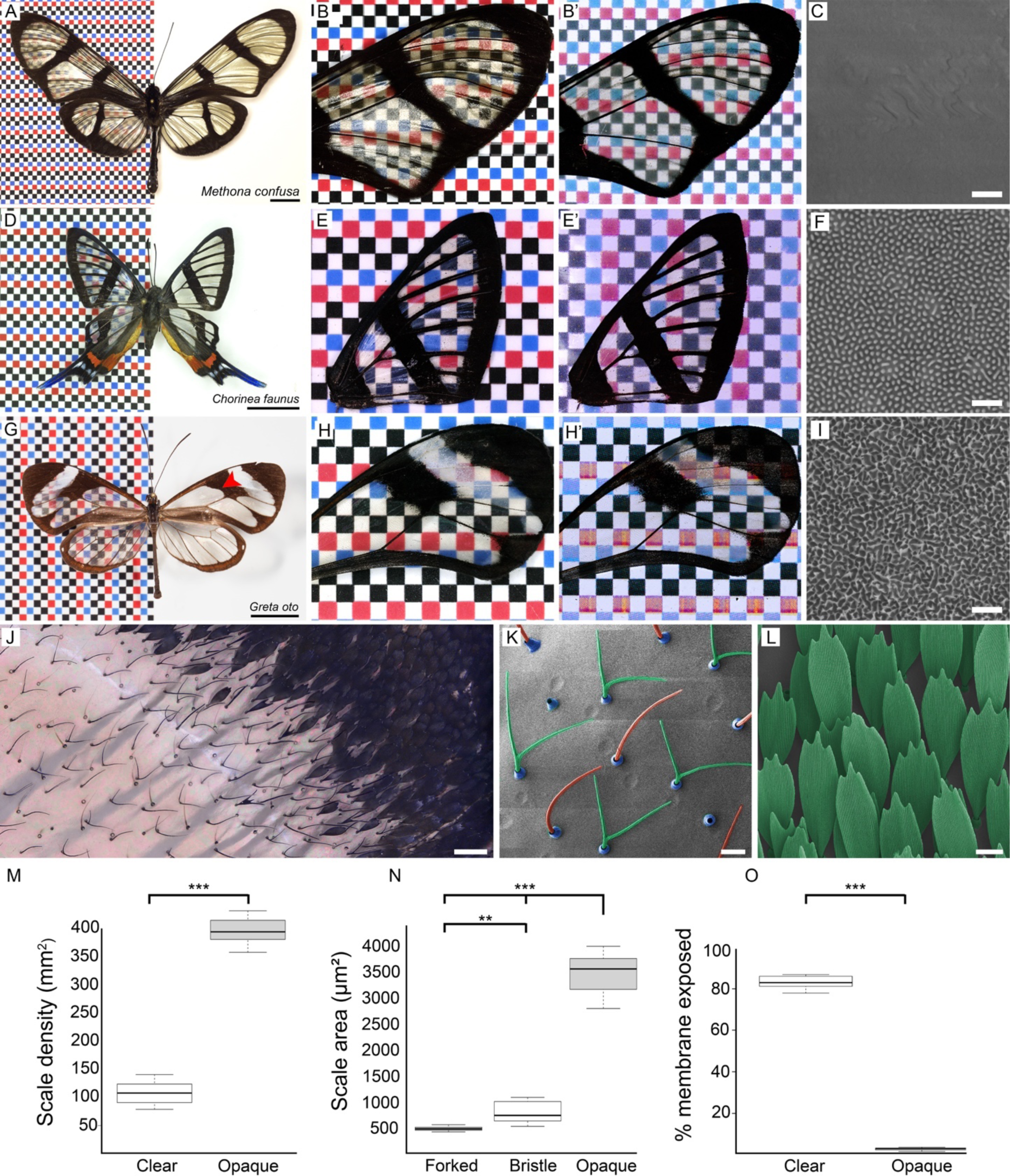
Examples of clearwing butterflies and wing scale features in *Greta oto*. **(A)** Giant glasswing *Methona confusa* (Nymphalidae: Ithomiini). Scale bar = 1 cm. Wings under **(B)** reflected and **(B’)** transmitted light illustrating general transparency, but strong light reflectance off the wing surface in this species. **(C)** The surface of the wing membrane is smooth and devoid of nanostructures. Scale bar = 1 μm. **(D)** Longtail glasswing *Chorinea faunus* (Riodinidae). Scale bar = 1 cm. Wings under **(E)** reflected and **(E’)** transmitted light illustrating minimal reflectance. **(F)** The membrane surface contains dome-shaped chitin nanoprotuberances that generate anti-reflective properties (*21*). Scale bar = 1 μm. **(G)** Glasswing *Greta oto* (Nymphalidae: Ithomiini). Red arrowhead indicates the representative clear and opaque wing region investigated, scale bar = 1 cm. Wings under **(H)** reflected and **(H’)** transmitted light illustrating minimal reflectance. **(I)** The surface of the wing membrane contains irregularly sized nanopillars that enable omnidirectional anti-reflective properties (*10*). Scale bar = 1 μm. **(J)** High magnification of a transition boundary between a clear (left side) and opaque (upper right side) wing region in *G. oto*. Scale bar = 100 μm. **(K)** SEM of adult scales in a clear wing region of *G. oto*, revealing the alternating forked (green false coloring) and bristle-like (red false coloring) scale morphologies (socket false colored in blue). Scale bar = 20 μm. **(L)** SEM of scales in an opaque wing region in *G. oto*, highlighting typical large, flat scale morphologies. Scale bar = 20 μm. **(M)** Measurements of scale density in clear and opaque wing regions, **(N)** scale surface area for forked, bristle-like, and opaque scale morphologies, and **(O)** percent of wing membrane exposed in *G. oto* clear and opaque regions. Error bars indicate means + SD of four measurements taken from wings in three different individuals, P-values are based on Student’s t-test for **(M), (O)**, and ANOVA test for **(N)**, ***P < 0.001; **P < 0.01.

Nevertheless, some organisms have evolved morphological innovations that overcome the challenges of terrestrial transparency, notably in the form of anti-reflective nanostructures. Early studies elucidated highly-ordered sub-wavelength nanostructures (termed ‘nipple arrays’) on the corneal surface of insect eyes (*17*). These structures were found to generally be ∼150-250 nm in height and spaced ∼200 nm apart, which reduces reflection across a broad range of wavelengths by creating a smoother gradient of refractive indices between air and chitin (*18*). Nanostructure arrays have also been identified on the wings of cicadas, which help to reduce surface reflection over the visible spectrum (*19*).

Some lepidopterans possess modified wing scales that allow light to reach the wing surface, which is composed of chitin and has some inherent transparency, but due to the high refractive index of chitin, *n* = 1.56 (*20*), the wing surface reflects light. For example, the butterfly *Methona confusa* (Nymphalidae: Ithomiini) has exposed wing membrane that lacks nanostructures on the surface, and as a result, the wing is somewhat transparent, but retains a high degree of reflectivity (Fig. 1A-C). Conversely, the longtail glasswing, *Chorinea faunus* (Riodinidae), contains small, widely spaced scales and dome-shaped chitin nanoprotuberances on the membrane that generate anti-reflective properties (Fig. 1D-F) (*21*). The hawkmoth, *Cephonodes hylas* (Sphingidae), has nude wings due to deciduous scales that fall out upon eclosion, and possesses anti-reflective nanostructures on its wing surface that morphologically resemble insect corneal nipple arrays (9). Nipple array nanostructures have also been characterized in transparent wing regions of the tiger moth *Cacostatia ossa* (Erebidae) (*22*). Finally, the glasswing butterfly *Greta oto* (Nymphalidae: Ithomiini) contains thin, vertically oriented scales, allowing the wing surface to be exposed, along with nanopillars that coat the surface. These irregularly arranged nanopillars feature a random height and width distribution and enable omnidirectional anti-reflective properties (Fig. 1G-I) (*10, 23*). More recent studies have explored aspects of structural diversity, optical properties, phylogenetic distribution, and ecological relevance of transparency within a wide range of butterflies and moths, highlighting that transparency has evolved multiple times independently and may present evolutionary benefits (*13, 24, 25*).

Lepidoptera are proving to represent an excellent group to investigate transparency on land, but the developmental processes underlying wing transparency are currently unknown. This presents a gap in our understanding of lepidopteran wing evolution and diversification, as transparent butterflies and moths contain multitudes of intriguing scale modifications and sub-wavelength cuticular nanostructures (*24, 25*). We therefore set out to explore the development of wing transparency in the glasswing butterfly *Greta oto*, which belongs to a diverse tribe (∼393 species) of predominantly transparent neotropical butterflies (*26*). We applied confocal and transmission electron microscopy to compare wing development, scale cytoskeletal organization, and membrane surface nanostructures between clear and opaque wing regions. Using chemical treatments, scanning electron microscopy, and gas chromatography–mass spectrometry, we found that nanostructures on the wing membrane surface are made of two layers: a lower layer of chitin-based nipple-like nanostructures, and an upper layer of wax-based nanopillars composed predominantly of long-chain *n*-alkanes. Finally, by removing the wax-based nanopillars, we demonstrate their role in dramatically reducing reflection on the wing surface via optical spectroscopy and analytical simulations.

## Results

### Scale measurements in clear and opaque wing regions of adult *Greta oto*

We investigated features of scale density, scale morphology, and the amount of wing surface exposed in wings of adult *Greta oto*. We focused on two adjacent regions within the forewing for consistency: a clear region within the discal cell and an opaque region that consists mainly of black scales near the M2-M3 crossvein. (Fig 1G,J). The clear wing region contained two types of alternating scale morphologies: bristle-like scales and narrow, forked scales, while within the opaque wing region, scale morphologies resembled ‘typical’ butterfly pigmented scales: flat and ovoid with serrations at the tips (Fig1. K,L). The mean density of scales (± SD) in the adult wing were significantly lower within the clear region (107 ± 19 scales per mm^2^) compared to the opaque region (395 ± 23 scales per mm^2^) (Student’s t-test, P < 0.001, n = 3 individuals, Fig. 1M). In the clear region, forked scales were significantly smaller in size (498 ± 39 μm^2^) compared to the bristle-like scales (831 ± 183 μm^2^), while in the opaque region, scales were the largest (3467 ± 382 μm^2^) (ANOVA test, n = 3 individuals, Fig. 1N). Finally, the amount of exposed wing membrane was significantly different between wing regions, with an average of 83.1% ± 0.76 and 2.4% ± 3.4 exposed membrane in the clear and opaque regions, respectively (Student’s t-test, P < 0.001, n = 3 individuals, Fig. 1O).

### Morphogenesis and cytoskeletal organization of developing scale cells

To investigate developmental processes of wing and scale development, we performed dissections of *G. oto* pupae at different time points (Fig. 2). As in other species of Lepidoptera, the early pupal wing consisted of a thin bilayer of uniform epithelial tissue and by 16 hours after pupal formation (APF) numerous epidermal cells had differentiated to produce sensory organ precursor (SOP) cells, which could be identified by fluorescently labelling tissue with DAPI (Fig. 2B,C) as the SOP’s are larger than, and positioned slightly basal to, the rest of the epidermal cells. The SOPs are precursors to the scale and socket cells and are organized into parallel rows. At this early stage of wing development, we observed that the clear wing region harbored a lower density of SOP cells relative to the opaque wing region (Fig. 2B,C). We can therefore infer that early into wing development, SOP cell patterning is differentially regulated between clear and opaque regions, which impacts the adult wing scale density and the amount of wing membrane surface exposed in different parts of the wing.

**Fig. 2.**
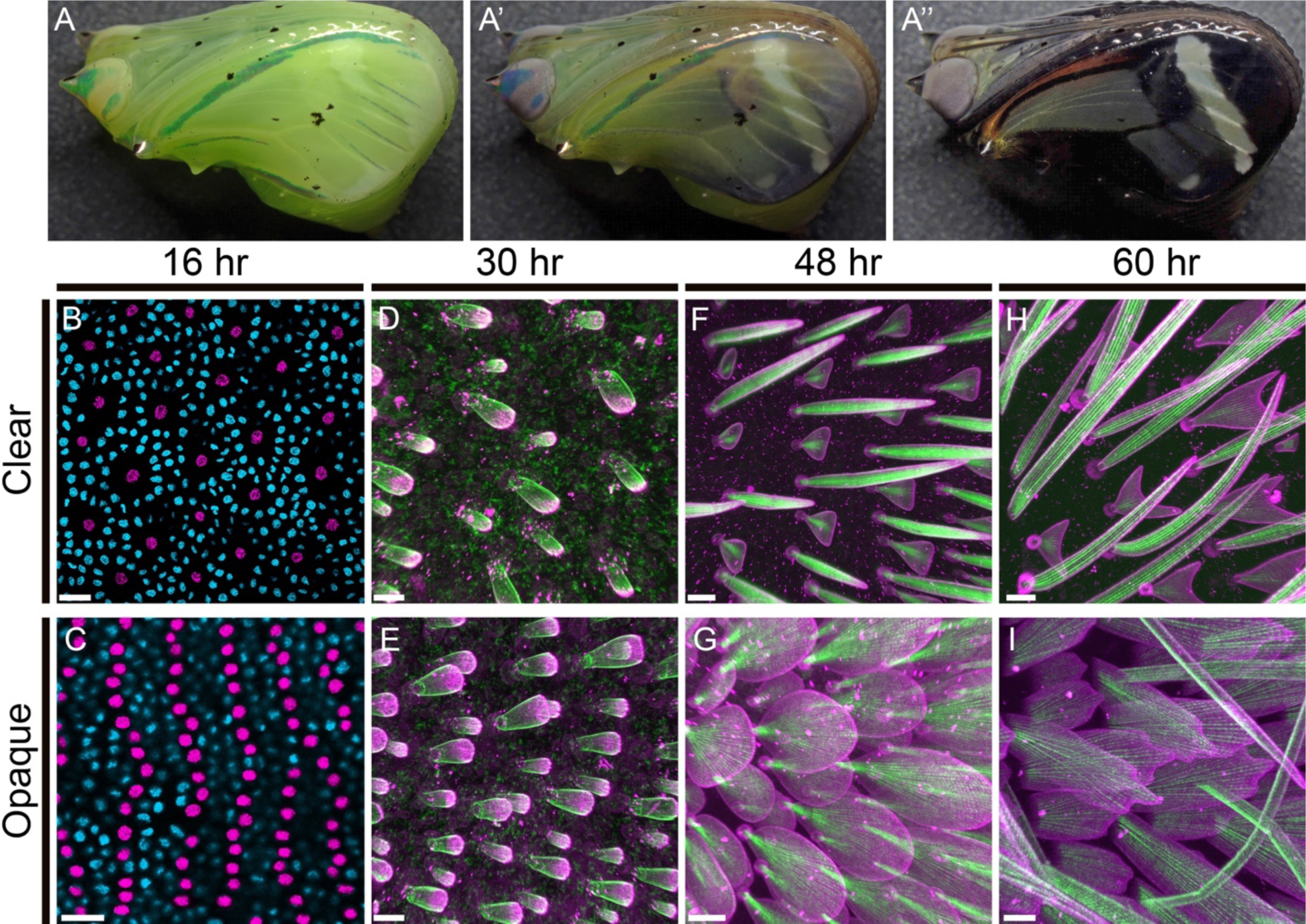
Pupal wing development and cytoskeletal organization of scales in clear and opaque regions. **(A)** Representative image of a *Greta oto* pupa ∼5 days after pupal formation (APF), **(A’-A’’)** developing up to the melanic stage ∼7 days APF, just prior to eclosion. **(B)** Early wing development 16 hours APF stained with DAPI (nuclei) in a clear wing region and **(C)** opaque wing region. The clear region contains a reduced number of sensory organ precursor (SOP) cells (which are the precursor cells to the scale and socket cells) relative to the opaque region. Scale bar = 20 μm. SOP cells are false colored magenta for better viewing. Simultaneous confocal imaging of fluorescently labeled scale cell membrane (wheat germ agglutinin; WGA, magenta), and F-actin (phalloidin, green), comparing clear wing regions **(D, F, H)** to opaque wing regions **(E, G, I)**. At 30 h AFP **(D, E)** WGA and phalloidin staining reveal early scale buds extending from the wing epithelium. F-actin reveals loosely organized parallel actin filaments protruding from the membrane. 48 h APF **(F, G)** scales have grown and changed in morphology. Short actin filaments have reorganized and formed smaller numbers of thick, regularly spaced parallel bundles under the surface of the cell membrane. **(F)** In the clear wing region, scale cells alternate between triangular shapes and bristles. 60 h APF **(H, I)**, developing scales have become more elongated. **(H)** The triangular-shaped scales in the clear wing region have proceeded to generate two new branches, which fork and elongate bidirectionally, while bristle-like scales have rapidly elongated and curved. **(I)** In the opaque region, scales are longer, wider, and have now developed serrations at the tips. Scale bar in **(D-I)** = 10 μm.

Next, we investigated cellular and cytoskeletal organization during scale growth in clear and opaque wing regions, using simultaneous confocal imaging of fluorescently labeled scale cell membrane (wheat germ agglutinin; WGA), and F-actin (phalloidin) (Fig. 2D-I). We found that general aspects of scale development in *G. oto* follow those previously reported in several butterfly and moth species by *(6)*, with some notable distinctions for modified scale growth in the clear wing regions of *G. oto*.

By 30 hours APF, the SOP cells have divided to produce the scale and socket cells (Fig. 2D,E). The scale cell body lies internally within the wing, while the socket cell associated with each scale cell lies in a more superficial position. At this pupal stage, the morphological development of wing scale projections has begun, and the scale cells develop as small buds containing short, densely packed parallel F-actin filaments. Phalloidin staining showed the appearance of these small cylindrical buds containing F-actin filaments, and WGA staining showed outlines of the membrane as the scale outgrowths begin to project and elongate beyond the wing surface. At this stage, budding scales in the clear wing region appeared morphologically similar to the unspecialized opaque scales: roughly elongated balloon-shaped with numerous small actin rods fanning out from the pedicel to the apical tip of the scale. In the clear region, early scale projections showed alternating sizes. In the opaque region similar budding scales at a higher density were found, with larger buds corresponding to future cover scales, and smaller, shorter buds corresponding to future ground scales (Fig. 2D,E).

By 48 hours APF, scale cell extensions have grown and elongated (Fig. 2F,G). The actin filaments have reorganized into smaller numbers of thick, regularly spaced bundles along the proximal–distal axis of the scale just under the surface of the cell membrane. At the base of the scales, fluorescent staining indicated that F-actin bundles are tightly packed, while in more distal regions we could see an asymmetric distribution of F-actin, with larger actin bundles in the adwing (facing the wing membrane) side of the scales (movie S1). At this stage, scales in different regions of the wing had also started to take on dramatically different morphologies. Scales in the clear region had elongated in a vertical orientation and obtained two types of alternating morphologies: short and triangular, or long and bristle-like outgrowths (Fig. 2F). The wings of other butterfly species contain alternating ground and cover scales, in which the ground scales are typically smaller in size than the cover scales, consistent with our observations of the opaque regions of *G. oto* (Fig. 2F). Based on scale size and position, we interpret that within the clear wing region of glasswing butterflies, the larger bristle-like scales are modified cover scales and smaller forked scales are modified ground scales (Fig. 2F). In the opaque region, scales have taken on a round and flattened morphology, similar to what has been described in other colorful butterfly and moth species, with the ground scales being shorter and wider than the cover scales (Fig. 2G).

By 60 hours APF, scale projections are even more elongated (Fig. 2H,I). The triangular scales in the clear wing region have proceeded to generate two new branches, which fork and elongate at the tips bidirectionally, while bristle-like scales have elongated and curved (Fig. 2H). In the opaque region, scales were longer, wider, flatter, and had developed serrations at the tips (Fig. 2I). F-actin bundles extended all the way to the distal tips of these serrations, which is necessary to produce finger-like projections at the tips of scales (*6*). Phalloidin staining also revealed that actin bundles were arranged in more symmetrical patterns around the periphery of the bristle-like scale morphologies, forked scales showed modified actin organization at the branching points, and actin bundle asymmetry was greatest in developing flat opaque scales, with larger bundles present on the adwing side (Fig. 2H,I).

### Ultrastructure analysis of developing bristle, forked and opaque scales

To reveal ultrastructural detail of developing wing scale morphology, we performed transmission electron microscopy (TEM) on pupal wing tissue of *G. oto* at 48 hours APF (Fig. 3). In transverse sections, we could resolve distinct scale morphologies (bristle, forked and opaque) and their associated cytoskeletal elements.

**Fig. 3.**
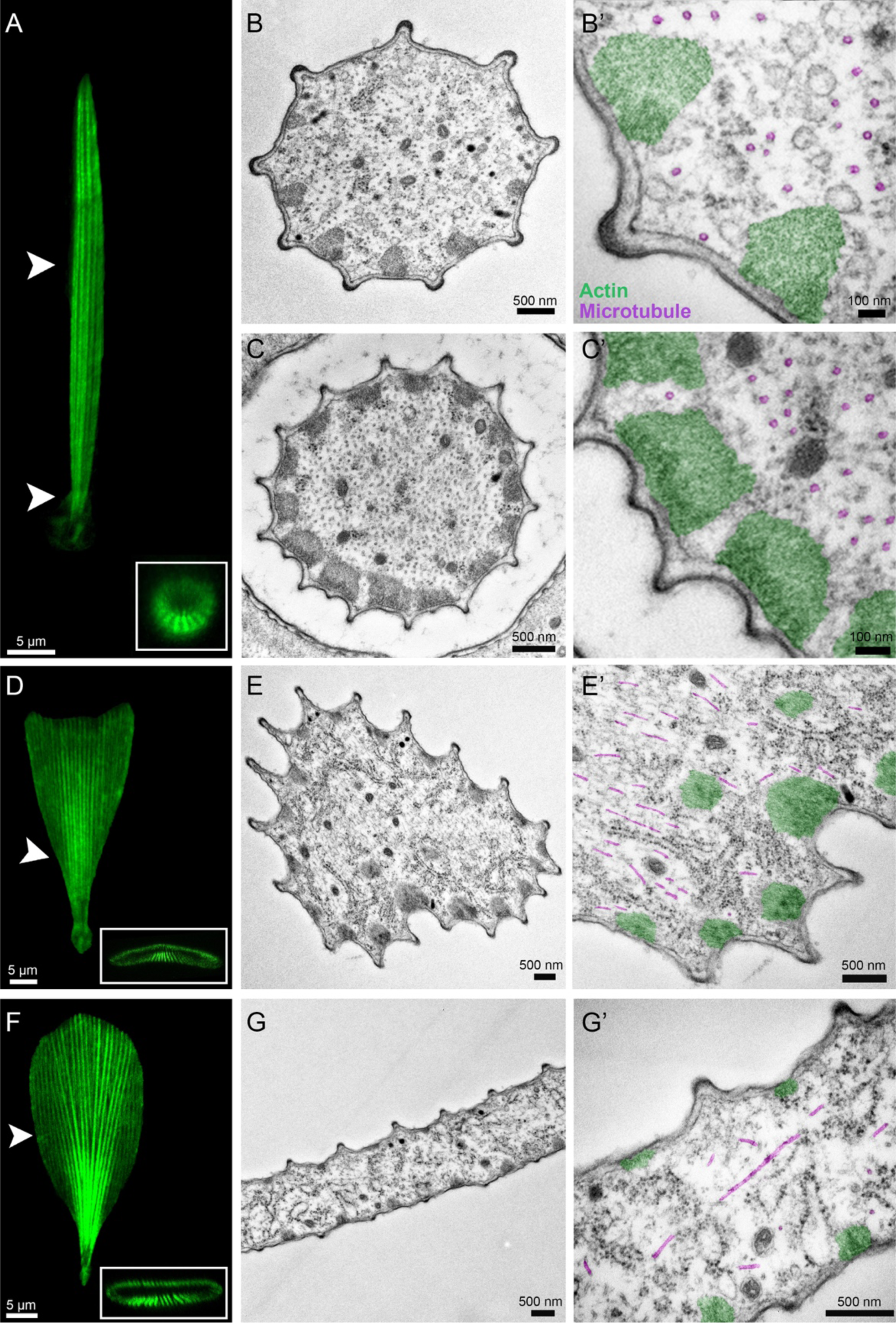
Confocal and transmission electron microscopy transverse sections of developing bristle (top), forked (middle) and flat (bottom) scales 48 hours APF. **(A)** Confocal projection of a bristle-like scale morphology (phalloidin) in a clear wing region. White arrowhead shows representative regions of transverse TEM sections. Scale bar = 5 μm. TEM of a bristle-like scale in a distal region **(B-B’)** and a basal region near the socket cell **(C-C’)**. Note the peripheral actin bundles (false colored green) and internal microtubule rings (false colored magenta). The more distal region of the scale **(B)** contains a lower density of microtubules relative to the base of the scale **(C)**. Scale bars in **(B**,**C)** = 500 nm and scale bars in **(B’**,**C’)** = 100 nm. **(D)** Confocal projection of a developing forked scale (phalloidin) in a clear wing region. White arrowhead shows representative regions of transverse TEM sections. Scale bar = 5 μm. **(E-E’)** TEM of a forked scale reveals peripheral bundles of actin (false colored green), with thicker actin bundles on the ventral side of the scale and internal microtubules (false colored magenta). Two internal bundles of actin filaments can be observed in the cytoplasm **(E’)**. Scale bars in **(E-E’)** = 500 nm. **(F)** Confocal projections of developing flat, round scale (phalloidin) in an opaque wing region. White arrowhead shows representative regions of transverse TEM sections. Scale bar = 5 μm. **(G-G’)** TEM reveals asymmetry in the actin bundles (false colored green), which are thicker on the bottom side of the scale relative to the upper surface. Microtubules (false colored magenta) are found in various orientations. Scale bar in **(G-G’)** = 500 nm.

Bristle-like scales in the clear wing regions were circular in cross sections (Fig. 3A-C). We could also distinguish between distal and basal regions of bristle-like scales, the latter of which had the presence of a surrounding socket cell in the cross section (Fig. 3B,C). TEM revealed that these bristle-like scales were ringed by peripheral bundles of actin filaments, which lay spaced just under the cell membrane (Fig. 3B-C’). On the adwing side of the scale, the actin bundles were larger and spaced closer to one another relative to the abwing side, and in more distal regions of the bristle-like scale, the actin bundles were more widely spaced and smaller in size. We also observed large populations of microtubules (MTs) distributed throughout the developing scales, which were internal relative to the actin bundles. Interestingly, we observed distinct patterns of microtubule distribution within different developing scale morphologies. The cross section of bristle-like scales revealed large populations of internal microtubules, which we identified due to their characteristic ring shape and diameter of ∼25 nm (Fig. 3B’,C’). The circular ring shape of microtubules in cross sections of both the basal and distal parts of the bristle-like scale suggested that microtubules are all longitudinally oriented, running in the same direction as the actin filaments, parallel to growth. We also observed that populations of MTs are localized primarily away from the surface of the scale in its interior, and MTs were fewer distally than basally (Fig. 3B’,C’).

In our TEM cross sections we also observed scale types that appeared more triangular in shape, suggesting that these corresponded to developing forked scales within the clear wing region (Fig. 3D,E). We observed that these scales were ringed by peripheral bundles of crosslinked actin filaments, with thicker actin bundles on the adwing side of the scale. Interestingly, we observed two internal bundles of actin filaments that were not observed in bristle-like scale morphologies (Fig. 3E’). We also note that there was variability in MT orientation, rather than the ubiquitous longitudinal orientations observed in bristle-like scales.

Finally, developing opaque scales were easily identified in cross sections due to their large size and flattened morphology (Fig. 3F,G). We observed peripheral bundles of crosslinked actin filaments that were widely spaced and smaller in size in distal parts of the scale (Fig. 3G-G’). We observed a clear asymmetry in actin bundle size, which were thicker on the adwing side of the scale relative to the abwing surface. In opaque wing regions, TEM micrographs revealed what appeared to be concentrated parallel-running populations of MTs near the narrow base of the scales, and then a more mesh-like network of MTs in more distal flattened regions, indicating that MTs have varying orientations within different regions of the scale (Fig. 3G,G’, fig. S1). In contrast to the bristle-like scales, large, flattened opaque scales appeared to contain populations of MTs that were more widely distributed and less dense.

In all scale types we observed the presence of numerous internal organelles and vesicles, including mitochondria, electron dense vesicles and free ribosomes (Fig. 3, fig. S1). We also observed that the actin bundles contained dense, hexagonally packed F-actin filaments, supporting previous patterns for actin bundle formation in elongating insect scales (fig. S1). The neck regions of different scale morphologies were predominantly filled with longitudinally oriented microtubules, actin bundles, and mitochondria. Longitudinal views also supported that MTs are numerous in the outgrowing scale, and their spatial arrangement differed with scale position and shape. More mature scales around 120 hours APF exhibited developed ridge morphologies and thickened cuticle layers (fig. S1).

### Ontogeny of wing membrane nanostructures

The clear wing regions of *G. oto* contain nanopillars that cover the surface of the membrane (Fig 1I, Fig 4A). These nanopillars were previously characterized in adult wings, which feature an irregular height distribution and help to generate omnidirectional anti-reflective properties (*10*). To gain insight into the development of these nanostructures, we examined the surface of the wing membrane epithelial cells with TEM (Fig. 4B-F). At 60 hours APF, a perpendicular section through the wing epithelia showed a continuous epithelial lamina (Fig. 4B,C). We observed the epithelial cells contained microvilli (MV), which appeared as slender linear extensions from the inner margins of the developing cells that insert into electron-dense material (Fig. 4B,C). The surface layer of the epithelia appeared as an extracellular lamellar system, and lamina evaginations appeared in the section as domes distal to the microvillar extensions (Fig. 4C). By 72 hours APF, we observed a thin outer layer of the epicuticle that rose above the epidermal cells and by 120 hours APF, we found that this upper layer above the microvilli contained what appear to be dome-shaped protrusions and thickened cuticle, possibly secreted from regularly spaced microvilli (Fig. 4D,E). Finally, in our TEM cross section of a fully developed adult wing of *G. oto*, we observed that the membrane surface harbors dome-shaped nanoprotrusions with similar morphologies to insect corneal surface nipple arrays (e.g. *9, 17*), which we refer to throughout the text now as “nipple nanostructures”, and an upper layer containing pillar-like protrusions, which we refer to as “nanopillars”, that featured a more irregular height distribution (Fig. 4F). These results show early subcellular processes of developing nanopillars within the clear wing region, which arise distal to microvillar extension in epithelial cells.

**Fig. 4.**
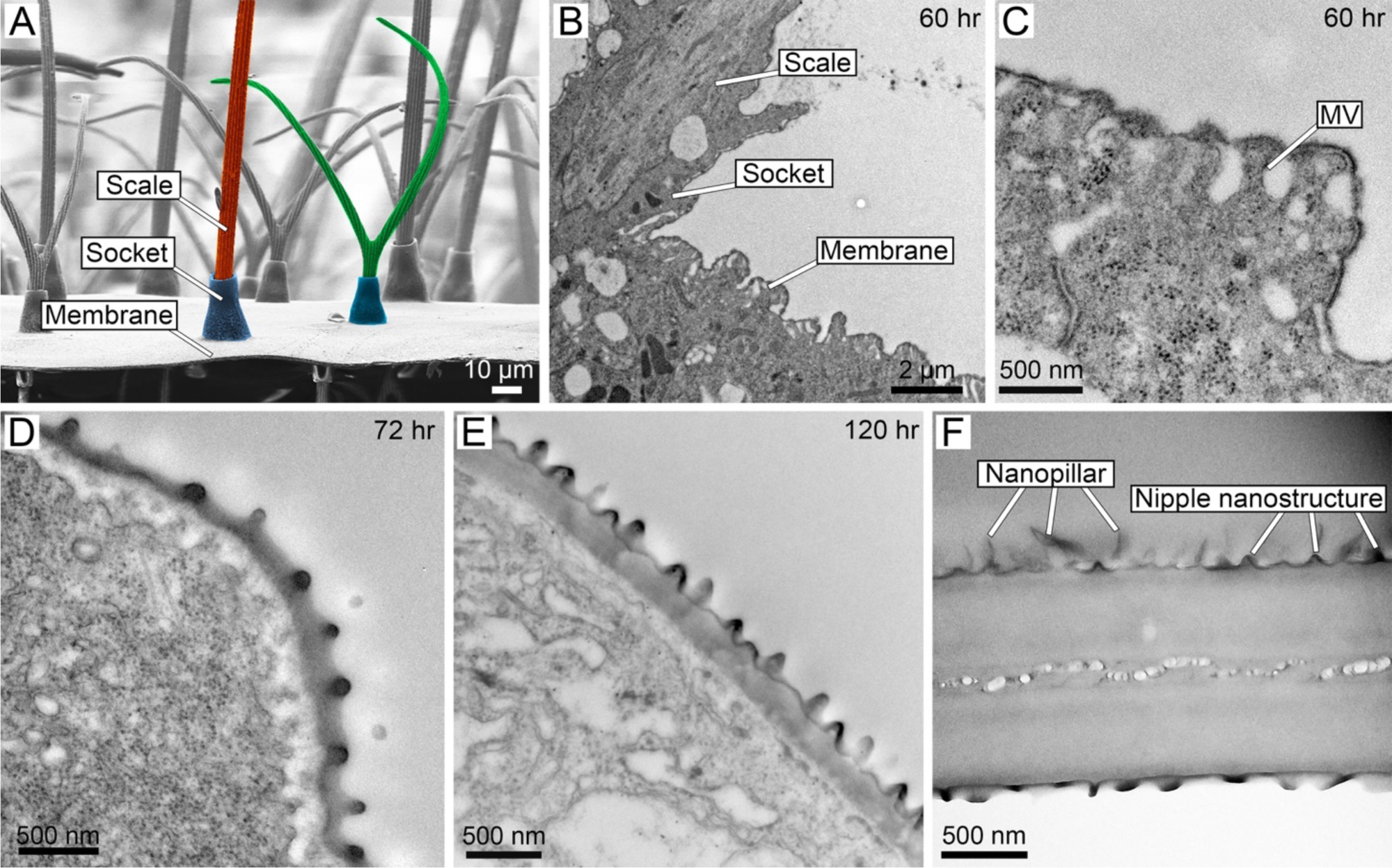
Ontogeny of wing membrane surface nanostructures. **(A)** SEM cross section (side view) of an adult *Greta oto* clear wing region. Scale bar = 10 μm. Bristle-like scale false colored in red, forked scale false colored in green, sockets false colored in blue. **(B)** TEM transverse section of epithelial tissue 60 h APF, showing lateral scale growth and wing membrane cells. Scale bar = 2 μm. **(C)** Higher magnification of developing wing epithelial cells at 60 h APF show microvilli (MV) projections, which appear as slender linear extensions from the inner margins of the developing cells that insert into a thin layer of electron-dense material. Lamina evaginations appear in the section as domes. **(D)** TEM of epithelial tissue 72 h APF and **(E)** 120 h APF shows wing surface nanostructures protruding from the surface, with tips of microvilli still attached to the inner surface of the wing membrane. **(F)** TEM of the adult wing membrane. The surface contains dome-shaped nipple nanostructures and an upper layer of nanopillars. Scale bar in **(C-E)** = 500 nm.

### Topographical organization and biochemical composition of wing surface nanostructures

Based on our EM results of membrane nanostructures, we investigated the topographical organization and biochemical composition of the adult wing surface. To do so, we treated individual, disarticulated adult *G. oto* wings in two ways: by 1) physically removing wing surface nanostructures by gently pressing and rubbing a wing in between paper and Styrofoam (after *9*) and 2) testing the wing surface structures for solubility in organic solvents, including hexane and chloroform to extract lipids (after *27*). We then performed SEM to compare wing surface topography of untreated and treated wing samples (Fig. 5A-C’). SEM confirmed that the first treatment partially or completely removed nanostructures across the wing membrane surface (Fig. 5B). In a region of partial removal, we could identify smaller, dome-shaped nipple nanostructures underneath the top layer of nanopillars (Fig. 5B’). SEM of the chemically treated wing surface revealed that the upper layer of irregularly sized nanopillars were completely removed, revealing a layer of regularly arranged dome-shaped nipple nanostructures that did not dissolve through chloroform or hexane exposure (Fig. 5C,C’). Therefore, we hypothesized that the upper layer of irregularly sized nanopillars consisted of a secreted wax-based material, which sits above smaller chitin-based nipple nanostructures.

**Fig. 5.**
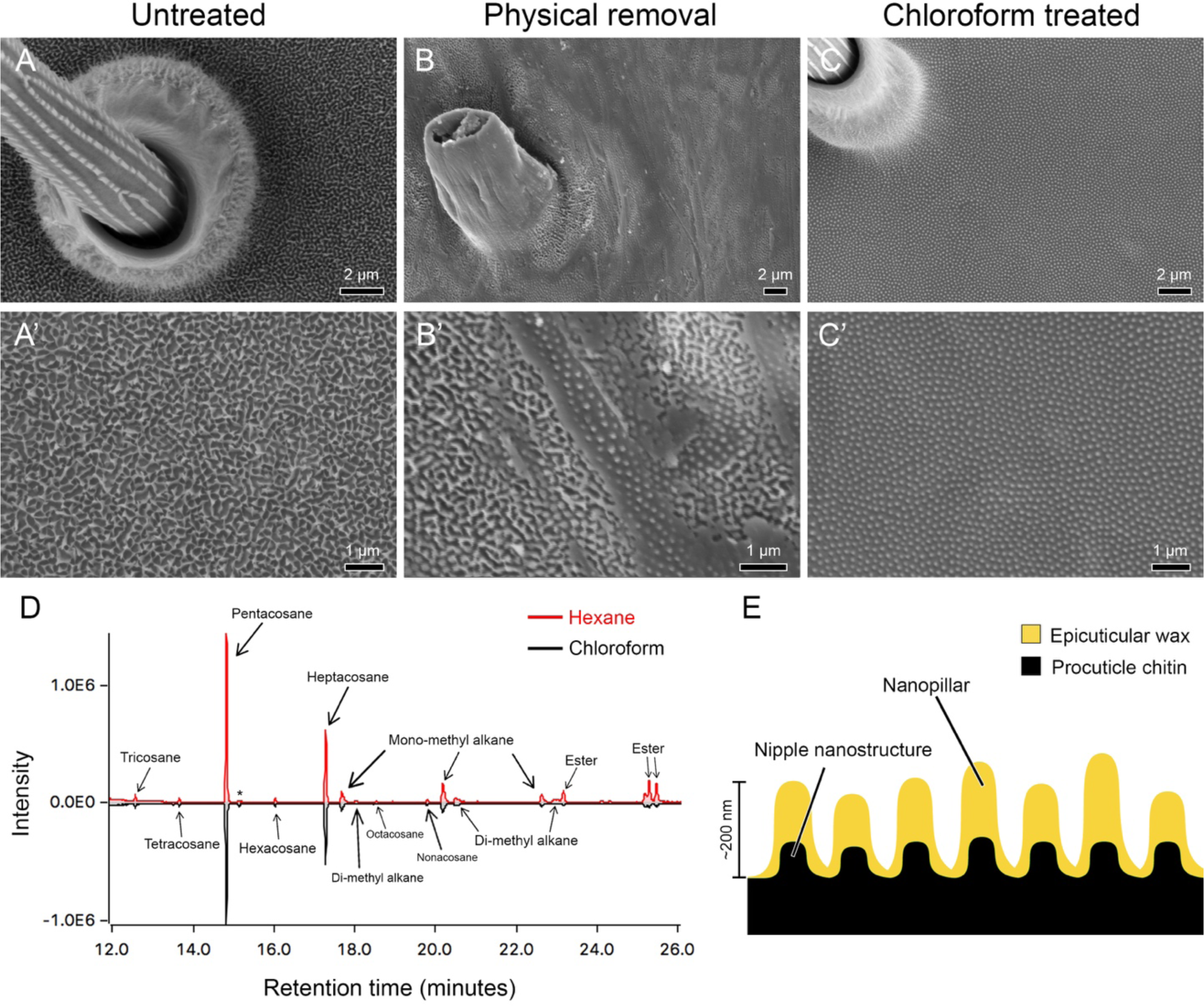
Topographical organization and biochemical composition of wing surface nanostructures. Scanning electron microscopy of the transparent wing membrane surface of *Greta oto* under **(A-A’)** untreated condition, highlighting the presence of irregularly arranged nanopillar structures covering the surface, **(B-B’)** physical treated condition, revealing partial removal of surface nanopillars, and a lower layer of more regularly arranged nipple-like nanostructures and **(C)** chloroform treated condition, revealing complete removal of the upper layer of nanopillars, and remining lower layer of nipple-like nanostructures. Scale bars in **(A, B C)** = 2 μm, scale bars in **(A’, B’, C’)** = 1 μm. **(D)** Chromatogram of hexane-treated (top; red line) and chloroform-treated (bottom; black line) clearwing extracts. X-axis shows the retention time in minutes and Y-axis shows the abundance of total ion current. **(E)** Schematic of proposed wing surface membrane nanostructures in *Greta oto*, composed of chitin-based procuticle and wax-based epicuticle.

To test this hypothesis, we extracted the surface layer of *G. oto* clear wing regions with either hexane or chloroform and analyzed the chemical composition by gas chromatography–mass spectrometry (GC-MS). We found that the chemical profile generated by both hexane and chloroform extracts yielded similar results (Fig. 5D). In all extracts, we identified two straight-chain alkanes that made up approximately 2/3 of the compounds detected: 41.64 ± 5.75% pentacosane (C_25_H_52_) and 23.32 ± 5.35% heptacosane (C_27_H_56_) (Table S1). The remaining compounds were primarily composed of slightly larger methyl-branched alkanes (monomethyl and dimethyl C27, C29 and C31) and esters. Therefore, our results suggest that in *G. oto* there are two components to wing surface ultrastructure: procuticle-based nipple nanostructures, and an upper epicuticular layer of irregularly sized nanopillars, composed mainly of straight chain alkanes (Fig.

5D,E).

### Anti-reflective properties of wax-based nanopillars

To address whether the wax-based nanopillars play a role in wing reflection, we measured the reflectance spectra of untreated and hexane-treated wings (Fig. 6). Additionally, we measured nanostructure geometries and membrane thickness from wing SEM cross sections (n = 6), and determined the average distance between two nanostructures as *d* = 174 nm, conical shaped cuticular nipple nanostructures height, *h*_p_ = 77 nm, wax-based irregular nanopillars radius, *r*_np_ = 53 nm, mean height, *h*_np_ = 224 nm and variance *σ*_np_ = 49.3 nm, and membrane thickness, *h*_m_ = 746 nm and variance *σ*_m_ = 43 nm (Fig. 6B,D, fig. S2). On the basis of SEM micrographs for treated and untreated samples, we modeled three wing architectures consisting of 1) nanopillars with variable height together with cuticle-based nipple nanostructures on the wing membrane, 2) cuticle-based nipple nanostructures on wing membrane and 3) wing membrane without any nanostructures, to simulate the optical properties for different conditions (Fig. 6E). The simulated reflectance data of the untreated and treated conditions in Fig. 6F closely resembled the experimental ones. In untreated wings of *G. oto*, we found that transparent regions have a low total diffuse reflection of about 2%, which is in line with previous reflectance measurements of this species (Fig. 6F, *10*). By contrast, the hexane treated wings without the upper layer of wax nanopillars had about 2.5 times greater reflectance relative to the untreated wings, and generated an iridescent thin film spectra, even though they harbored dome-shaped nipple nanostructures (Fig. 6D,F).

**Fig. 6.**
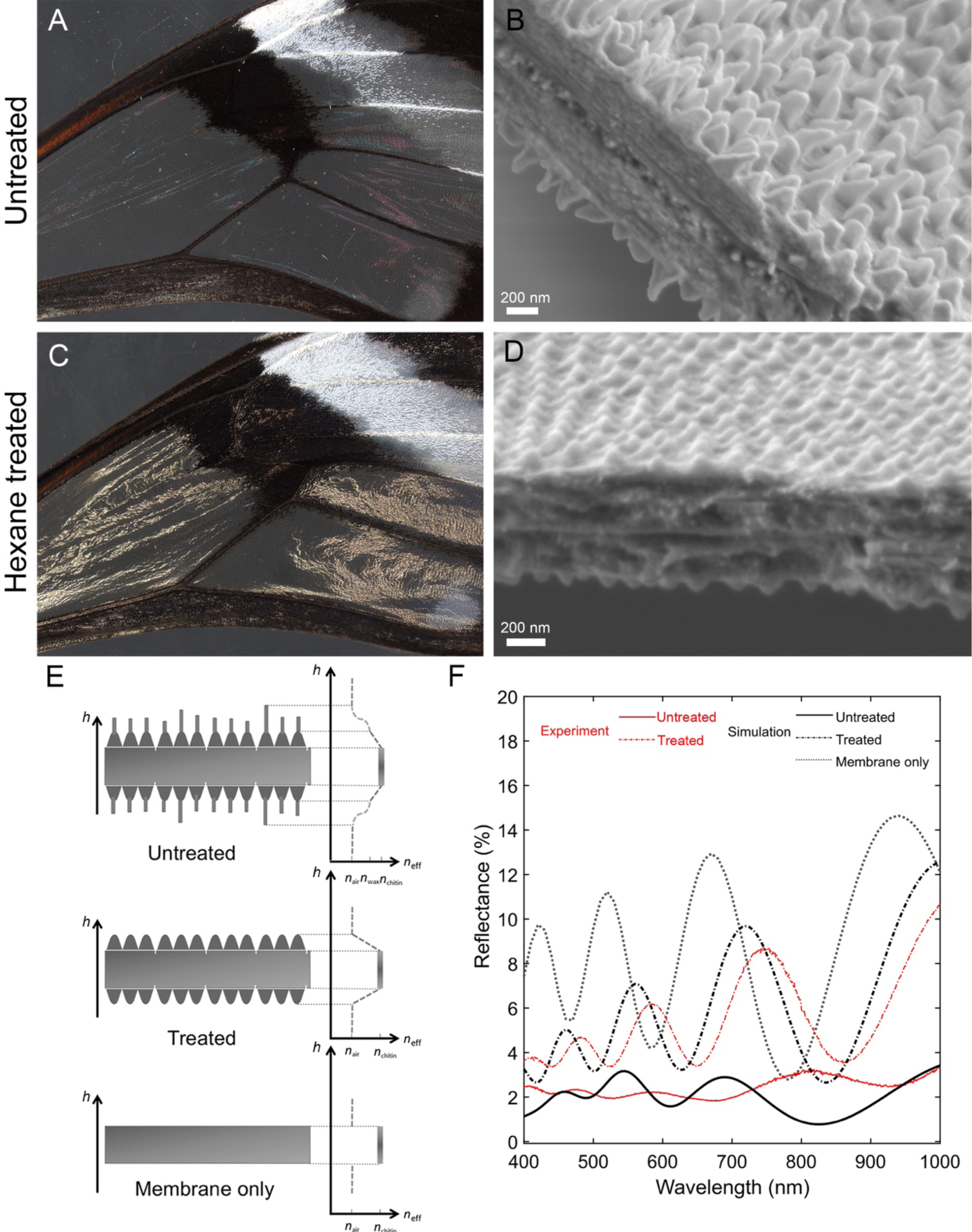
Structural elements, reflectance spectra and optical modeling of anti-reflective nanostructures. Optical images and cross section scanning electron microscopy of *Greta oto* **(A, B)** untreated wings, illustrating low reflectance and the presence of nanopillars on the wing membrane surface and **(C, D)** hexane-treated wings illustrating increased reflectance and the loss of nanopillars on the wing membrane, but presence of nipple-like nanostructures on the surface. Scale bars for **(B, D)** = 200 nm. **(E)** Optical modeling of effective refractive index conditions for untreated (top), with nanopillars of variable height together with cuticle-based nipple nanostructures on the wing membrane, treated (middle) with cuticle-based nipple nanostructures on wing membrane and wing membrane without any nanostructures (bottom). Y-axis represents height *h* and X-axis represents effective refractive index condition of air (n_air_), chitin (n_chitin_), and wax (n_wax_). **(F)** Representative reflectance spectra of experimental (red) and simulation data (black) for untreated wings with nanopillars on the membrane surface (solid line), hexane-treated wings with the wax-based layer of nanopillars removed (dashed line) and membrane only (dotted line).

For simulated data, the overall reflectance ratio of the hexane treated wing to that of the untreated was approximately three, similar to experimental reflectance data (Fig. 6F, Table S2). Most importantly, the simulated results for the untreated wing with wax-based irregular nanopillars make reflectance more uniform across wavelengths, which reduces the iridescent effect of the wing membrane. Finally, we simulated a thin film membrane without any nanostructures, which showed reflectance (averaged from all wavelengths) of the membrane itself to be 8.81 ± 3.46%, whereas the treated and untreated wing reflections were 5.78 ± 2.82% and 1.93 ± 0.77%, respectively (Fig. 6F). While treated wings harboring dome-shaped nipple nanostructures reduced the overall reflectance relative to the membrane only, their effect was not strong enough to reduce reflectance spectra oscillation. The wax-based irregular nanopillars on top introduced a more gradual transition between refractive indices to lessen the oscillation by approximately five-fold, in addition to reducing overall reflection (Fig, 6F). Additionally, we simulated the three wing architecture models considering different mean membrane thicknesses and variance in membrane thickness (fig. S3). We found that variance in wing membrane thickness reduced reflectance spectra oscillations, rather than mean membrane thickness alone, and more peaks appear in the visible spectrum with increasing thickness of the membrane. (fig. S3, Table S3). Overall, these results demonstrate that the non-constant architecture of the wing membrane and wax-based irregular nanopillars on the wing surface of *G. oto* function to dramatically enhance anti-reflective properties.

## Discussion

Butterflies and moths have evolved sub-wavelength anti-reflective structural innovations on their wings that enable them to be transparent. Here we report the details of pupal wing development and cytoskeletal organization in the glasswing butterfly, *Greta oto*, as well as insights into the ontogeny and biochemical basis of wing surface nanostructures that reduce reflection.

The arrangement of unicellular projections in insect integument, such as bristles and scales, has been a model for research on cellular pattern formation (*28*). Shortly after pupation, sensory organ precursor (SOP) cells develop from a monolayer of epithelial cells into orderly arrangements, then differentiate into scale and socket cells. In the present study, we found that early SOP cell patterning impacts the final adult scale density in *G. oto* and this feature of spacing scale cells farther apart, and therefore reducing the overall density of scales, is an initial step to generate clear wings. During early pupal development, the receptor molecule Notch is expressed in a grid-like pattern in the wing epithelium (*29*). This may contribute to the parallel rows of uniformly spaced SOP cells that express low levels of Notch, likely through a lateral inhibition mechanism. The low-Notch SOP cells express a homolog of the *achaete-scute* proneural transcription factors, which likely plays a role in scale precursor cell differentiation (*30*). Notch-mediated lateral inhibition could establish a dense population of ordered SOP cells in the developing wing, resulting in a characteristic ratio of scale-building and epithelial cells. Future studies should investigate if modifications in Notch signaling play a role in scale cell patterning in clearwing butterflies and moths, many of which contain reduced densities of scale cells (*24, 25*).

The range of morphological diversity among scales and bristles within Lepidoptera likely results developmentally from components or modifiers of the cytoskeletal structures and cell membrane. One study surveyed a wide range of developing butterfly and moth scales and identified that F-actin is required for several aspects of scale development, including scale cell elongation and proper orientation (*6*). In the present study, we found that *G. oto* serves as an excellent model to study differences in bristle and scale morphogenesis, as the wing contains a wide range of different scale types. In the developing bristle-like scales, we find symmetrical actin bundles that outline the cell periphery and a large population of longitudinally running interior microtubules. This is similar to what has been described for developing bristles in *Drosophila melanogaster* pupae, which contain peripheral bundles of cross-linked actin filaments and a large population of microtubules that run longitudinally along the bristle (*31*). Recently, (*32*) showed that actin bundles play different roles in shaping scales and bristles in the mosquito *Aedes aegypti*, in which developing bristles contained symmetrically organized actin bundles, while actin bundle distribution in scales became more asymmetrically organized. Given that actin dynamics play a variety of roles in regulating the development of bristles and scales (*6, 7, 32, 33*), we hypothesize that modifications in F-actin organization of scales in the transparent wing of *G. oto* are responsible in part for their narrow bristle-like and forked morphologies. In *D. melanogaster*, subunits of actin are rapidly added to the barbed ends of the actin filaments of bristles, relying on actin polymerization and bundling for this purpose, and cross-linking proteins are required early to bring filaments together (*31*). One cross-linking protein, Fascin, connects filaments together into hexagonally packed bundles. Our TEM of actin bundles, along with previous studies, support a similar mechanism of hexagonally packed F-actin bundles in Lepidoptera (Fig. 3, fig. S1) (*6, 7*).

In animal cells, microtubules have been frequently observed in arrangements parallel to the long axis of cellular extensions, such as axons, dendrites, and developing lepidopteran scales (*33*). In an analysis of moth scale development, major shape changes were found to be correlated with changes to the orientation of the cytoplasmic microtubules (*33*). In the present study, we identified large populations of microtubules organized throughout developing scales and showed that microtubules are more concentrated at the base of the scale. We also found that microtubules exhibit different distributions and orientations relative to distinct scale morphologies, namely between bristle, forked, and flat, round scales. In *D. melanogaster*, it has been suggested that bristle microtubules play a role in elongation, noting that they are highly stable, form at the start of the elongation, and then extend along the shaft as the cell elongates (*30*). A more recent reinvestigation of the role of MTs in *D. melanogaster* bristle elongation suggests that two populations of microtubules help to guide bristle development: dynamic microtubules (with mixed polarity) add bulk to the bristle cytoplasm and are thought to contribute proper axial growth, while stable microtubules act to polarize the axis of bristle elongation and are believed to aid organelle and protein distribution (*34, 35*). It would be interesting for future studies to functionally characterize the role microtubules play in the development of lepidopteran scales. Overall, we found conservation of developmental processes in scale formation relative to other previously described Lepidoptera, with notable differences in clear versus opaque wing regions. These findings lend further support that general patterns of scale development, including patterns of F-actin localization and microtubule distribution, seem to be well conserved in Lepidoptera, and that modifications of scale morphology to achieve clearwing phenotypes, such as narrow bristle-like and forked scales, likely involve alteration of cytoskeletal organization during scale growth.

Chitinous wing membrane has a higher refractive index than air, so as a mechanism that reduces glare, some clearwing species have evolved sub-wavelength anti-reflective nanostructures (*9, 10*). In this study, we identified the early developmental processes of nanostructures that arise in the wing epithelium. We also note interesting parallels of our observations to previous descriptions of developing nanostructures on the surface of insect cornea. Early data on pupal development of corneal nanostructures were produced by detailed electron microscopy studies, showing that corneal nipples emerge during lens formation, and a chitinous layer may be subsequently secreted underlying the nanostructure (*36*). In these observations, development of initial laminar patches formed on top of underlying microvilli. Subsequently, nanostructures (termed nipple structure array) formed on the surface, with the tips of microvilli still attached to the inner surface. Another study subsequently investigated pupal eye development in *D. melanogaster* and identified features of corneal nipple array formation that matched observations previously made in moth eye nanostructure development (*37*). Gemne (*36*) proposed that the corneal nanostructures originate from secretion by the regularly spaced microvilli of the cone lens cells, although there is still debate about the exact nature of how microvilli pre-pattern nanostructure arrays (*38*). Our TEM results provide insight into the early developmental processes of anti-reflective nanostructure formation in the wings of *G. oto*, highlighting certain similarities to nipple array development in insect cornea. It would be interesting for future work to explore if features of nanostructure formation arose independently in insect cuticle as a mechanism to reduce surface reflection.

In contrast to previously described highly ordered nipple arrays on insect eyes (e.g. *18, 38*), the irregularly sized anti-reflective nanopillars in the clear regions of *G. oto* wings consist of an upper layer of wax-based epicuticle sitting above procuticle-based nipple nanostructures. Insect cuticle is an extracellular matrix formed by the epidermis and composed of three layers: the outermost envelope, the middle epicuticle and the inner procuticle (*39*). The envelope and the epicuticle are composed mainly of lipids and proteins, while the procuticle contains the polysaccharide chitin. Many terrestrial arthropods deposit a layer of wax lipids on the surface of their cuticle, which reduces evaporative water loss (*40*). In some species of dragonfly, epicuticular wax-based nanostructures have also been demonstrated to play a role in generating optical properties, such as an ultraviolet reflection (*27*). In mature males of the dragonflies, a dense wax secretion composed of long-chain methyl ketones, in particular 2-pentacosanone, was found to contribute to the UV reflection properties. The chemical composition of nanopillars on the wing surface of cicadas, which have been shown to contribute to wettability and antimicrobial properties, and found that the major epicuticular components are fatty acids and hydrocarbons ranging from C_17_ to C_44_ (*41*). Another study exploring the molecular organization of dragonfly wing epicuticle found that the major components identified were fatty acids and *n*-alkanes with even numbered carbon chains ranging from C_14_ to C_30_ (*42*). Here, we identified that the epicuticular layer of irregularly sized anti-reflective nanopillars in *G. oto* appear to be composed mainly of *n*-alkanes, including pentacosane (C_25_) and heptacosane (C_27_) and showed the importance of these structures to attain better transparency.

Due to thin film optics, the thin membranes of insect wings sometimes reflect distinct structural coloration and iridescence (*43*). However, variability and non-constant thickness render the wing membranes as non-ideal thin films, and additional surface nanoprotrusions can introduce a gradient of refractive indices that reduces thin film reflections (*44*). For instance, membrane thickness was found to vary over the transparent wings of the damselfy *Hetaerina americana* from below 1 μm to up to 3 μm, yet membrane nanoprotrusions acted as an effective impedance matching device to reduce reflectance (*44*). In that study, average reflectance spectra for the Andromica clearwing butterfly *Greta andromica* was also calculated, although the wing was treated as a thin film, and did not address membrane surface nanostructures. By varying thickness in a Gaussian way while maintaining average thickness, (*44*) found that an increasing width of the Gaussian progressively reduced modulation of the reflectance spectrum. Similarly, in the present study, measurements from SEM cross sections of *G. oto* transparent wings indicate that the membrane thickness is non-constant, and in our optical simulations, variance in membrane thickness was found to be an important parameter for reduced reflectance spectra modulation (fig. S3). Overall, we found that variance in membrane thickness and wax-based nanostructures with irregular height distributions in *G. oto* reduce iridescence and maintain anti-reflection properties, which likely aid in crypsis (*11*).

Turing reaction-diffusion mechanisms have been proposed as a model for the formation of various corneal nanostructure morphologies (such as spacing, height, and spatial organization) during insect eye development (reviewed in *38*). Although the degree of height irregularity of nanopillars is important for achieving omnidirectional anti-reflection in *G. oto*, we do not yet understand how the wax nanopillars are generated to vary in height. Perhaps the pressure of the wax secretion varies across the microvillar extensions’ area, similar to how nozzle area plays a role in the propulsion force, and tunes the height of the nanopillars in the process. In such a scenario, the degree of the height variation could be synthetically engineered depending on the two-dimensional nanopatterned mask design in the biomimetic processes, like molding or imprinting techniques. Additionally, others have generated three-dimensional wax structures by using *n*-alkanes, noting that wax-based crystals can generate different shapes, sizes and densities depending on the chain length (*45*). Future work should investigate the possible role of alkanes, and the two-dimensional surface growth geometry, in generating three-dimensional anti-reflective nanostructures and potential applications for biomimetics. Our exploration of *Greta oto* wing development can serve as a model for understanding how transparent phenotypes evolved within Ithomiini, a diverse tribe of neotropical butterflies that act as mimicry models for numerous species of Lepidoptera (*26*), as well as more distantly related butterfly and moth species.

## Materials and Methods

### Samples

Glasswing butterfly (*Greta oto*) pre-pupae were purchased from Magic Wings Butterfly House (Deerfield, Massachusetts, USA) and reared on *Cestrum nocturnum* (Solanaceae) leaves at 27°C and 60% humidity on a 16:8 hour light:dark cycle at the Marine Biological Laboratory (Woods Hole, MA) under the United States Department of Agriculture permit number P526P-19-02269. At the appropriate time of development, pupal wings were dissected and age was recorded as hours after pupal case formation (h APF) as in (*6*). The average timeline from pupation to eclosion (adult emergence) for *G. oto* at 27°C is about 7 days, and we report our time series here which covers early aspects of wing scale development.

### Optical imaging and scale measurements

Images of whole mounted specimens were taken with a Canon EOS 70D digital camera with an EF 100mm f/2.8L macro lens. High-magnification images of disarticulated wings were taken with a Keyence VHX-5000 digital microscope. Scale density was determined by counting the numbers of scales in a 1 mm^2^ area. Scales were also removed from the wings, laid flat onto a slide, and Keyence software was used to measure the surface area of individual scales. Images of clear and opaque regions were processed with Keyence software to measure the percentage of area covered by scales. Sample size was equal to three individual butterflies reared in the same cohort, in which four measurements for each individual were averaged. We performed Student’s t-tests for scale density and percent of exposed membrane, and one-way ANOVA test for scale surface area comparisons.

### Confocal microscopy

For confocal microscopy of fixed tissue, pupal wings were dissected and fixed in PIPES, EGTA, MgSO_4_ (PEM) buffer with 3.7% paraformaldehyde for 20-30 minutes at room temperature, as described previously (*6*). Fixed wings were incubated in 1X PBS+0.1% Triton-X 100 (PT) with 1:200 dilution of phalloidin, Alexa 555 conjugated (Invitrogen A34055), and Wheat Germ Agglutinin, Alexa 647 conjugated (Invitrogen W32466) at a dilution of 1:200 overnight at 4°C. Wings were washed in PT and then placed in 50% glycerol:PBS with 1 µg/mL DAPI overnight at 4°C. Wing samples were placed on microscope slides and mounted in 70% glycerol:PBS. A coverslip (#1.5 thickness) was applied, and each preparation was sealed with nail polish. Slides of fixed tissue were examined with an LSM 880 confocal microscope (Carl Zeiss, Germany) with 40x and 63x objectives. Confocal images and movies were generated using Imaris Image Analysis Software (Bitplane, Oxford Instruments, UK).

### Scanning electron microscopy

We cut 2mm square pieces from dry wings, coated them with a 10 nm layer of gold using the BIO-RAD E5400 Sputter Coater, and imaged with a Hitachi TM-1000 SEM at 5 kV. Top-view and cross section SEM images were analysed with ImageJ 1.52 to measure membrane thickness and nanostructure dimensions (n = 6).

### Transmission electron microscopy

For transmission electron microscopy, wings of *Greta oto* pupae were dissected and fixed in 2% glutaraldehyde, 2% paraformaldehyde in 0.1 M sodium cacodylate buffer overnight at 4°C (pH 7.4). Samples were then rinsed in 0.1 M cacodylate buffer (pH 7.4) and post-fixed in1% aqueous osmium tetroxide in 0.1M cacodylic buffer overnight at 4°C, then rinsed in water. Samples were en bloc stained with 1% uranyl acetate in water and then rinsed in water. Samples were dehydrated through a graded ethanol series (50–100% in 10% steps), rinsed in propylene oxide, then infiltrated in 50% resin and propylene oxide overnight. Samples were infiltrated with Epon/Alardite embedding medium (70%, 80%, 95% to 100% steps) and polymerized at 60°C for two days. Thin sections (∼70nm) were cut on an Ultramicrotome RMC PowerTome XL using a Diatome diamond knife. Digital images were taken using a JEOL 200 transmission electron microscope (Jeol, USA).

### Wing surface wax extraction and analysis

To identify the molecular composition of the transparent wing surface, we pooled wing dissections from three individual adults and performed two replicates for chloroform-based extractions and two replicates for hexane-based extractions (after *26*). First, the samples were soaked with 100 µL of either hexane or chloroform and gently mixed for 15 minutes on a Thermolyne RotoMix 51300. The liquid solutions containing dissolved wing surface compounds were then transferred to glass vials with fixed microvolume inserts and the solvent was evaporated under a stream of high-purity nitrogen gas (99.99%). Dried extracts were re-dissolved in fixed volumes of hexane (10 µL), and half of the extract (5 μl) was injected by automatic liquid sampler into a gas chromatograph coupled with a mass selective detector (GC: 7890A; MS: 5975C; Agilent Technologies, USA) operating in electron impact mode. The injection was performed in a split/splitless injector in the splitless mode. Separation of compounds was performed on a fused silica capillary column (DB-5MS, 30 m × 0.32 mm × 0.25 μm, Agilent J&W GC columns, USA) with a temperature program starting from 80°C for 5 min and increasing by 80°C per min to 200 °C, followed by an increase of 5 °C per min to 325 °C which was held for 3 min, with helium used as the carrier gas, positive electron ionization (70 eV), Analog to Digital (A/D) sampling rate was set at 4, and the scan range was m/z 40.0 to 650.0. Chemical data processing was carried out using the software “Enhanced Chemstation” (Agilent Technologies, USA). We retained peaks with abundances greater than 0.25% of the total and compounds were identified according to their retention indices, diagnostic ions, and mass spectra, which are provided in Table S1. For some peaks, it was not possible to narrow the identity to a single specific compound because (1) some low abundance substances produced poor quality mass spectra, (2) multiple compounds could have produced the observed fragmentation patterns and/or (3) multiple compounds may have co-eluted at the same retention time.

### Optical measurements

The wing reflection measurements were performed on a Cary 5000 UV-Vis-NIR spectrophotometer, equipped with a light source of tungsten halogen and an integrating sphere diffuse reflectance accessory (Internal DRA 1800). Wing measurements from the dorsal wing surface (n = 6) were recorded with unpolarized light with a spot size of 100 µm for an incident angle of 8° to avoid the loss of direct specular reflectance component through the aperture. All measurements were taken in the dark to avoid possible stray illumination from the surrounding environment. A reference measurement was done with a calibrated commercial white spectralon standard to calculate the relative diffuse reflectance. The reflectance measurements and mean data are presented in Table S2.

### Optical simulations

The reflectance of the wing membrane before and after chemical treatment by hexane was analytically modeled using effective medium theory and transfer matrix method (*10, 18*). First, the effective volume fraction of the nanoprotuberances before and after the chemical treatment were based on measurements taken from SEM micrographs of the wings. We used the average distance between two hexagonally arranged nanostructures, *d*, conical shaped nipple nanostructures with height, *h*_p_, wax-based irregular nanopillars with radius, *r*_np_, mean height, *h*_np_ and variance *σ*_np_, and membrane thickness, *h*_m_ and variance *σ*_m_ (fig. S2). We considered a Gaussian distribution of irregular nanopillar height, as described previously (*10*). We also modeled the membrane thickness with Gaussian distribution to replicate the experimental membrane modulation in the calculation (*44, 46*). The total volume fraction of the untreated wing along the height *h* can be given by:

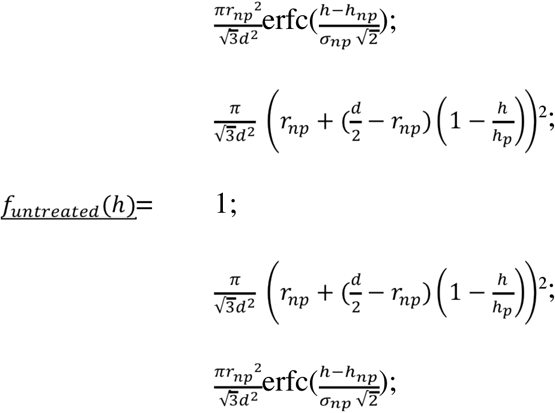

where, 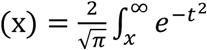 is the complementary error function.

The volume fraction of the treated wing without the irregular nanopillars will be:

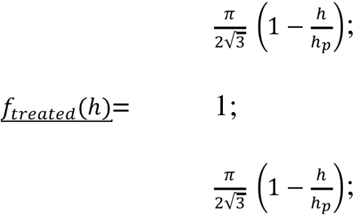

After determining the volume fraction, the corresponding refractive index changes along the wing at any height *h* was calculated using the effective medium theory with the Maxwell-Garnett approximation as shown in Fig. 6E, fig. S2. The refractive indices of the different materials were considered as n_air_ = 1, n_chitin_ = 1.56 + i0.008 (*20, 21*) and we considered n_wax_ = 1.39 (based on *47*). Afterwards, the transfer matrix method computed the reflectance from the stratified medium with calculated refractive index profiles as shown in Fig. 6E for the unpolarized condition (taking the average of both s- and p-polarization) at an incident angle of 8°. The membrane-only reflection at normal incident light can be directly calculated from (*46*):

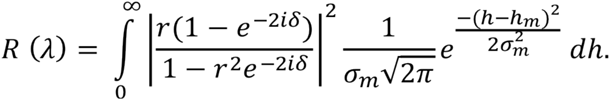

Where, *δ* = (2*πn*_chitin_*h*)/*λ* is the phase delay introduced by the membrane thickness of *h*, and *r* is the reflection coefficient at the air-chitin boundary governed by Fresnel’s equation for a normal incident light, i.e., *r* = (1 - *n*_chitin_) / (1 + *n*_chitin_).

## Acknowledgments

We would like to thank the Angie Serrano, Paola Betucci, Idoia Quintana-Urzainqui and Helena Bilandzija from the MBL Embryology Course and Cao Lu Yan from the MBL Physiology Course for their preliminary work on scale morphology of clearwing Lepidoptera, and subsequent work by Jaap van Krugten, Raymundo Picos, and Johnny On at UC Berkeley. We also thank Rachel Thayer, Arnaud Martin, Damien Gailly, Melanie Mcclure, Luca Livraghi, Oscar Paneso, Rémi Mauxion and Owen McMillan for their assistance with fieldwork, rearing, and preliminary experiments. We thank Fred Gagnon of Magic Wings Butterfly Conservatory and Gardens for assistance with butterfly rearing and Neil Tsutsui for support with the GC-MS experiments. Members of the Patel Lab, Craig Miller, and Noah Whiteman provided helpful feedback on the manuscript. RHS acknowledges the support from the Beckman Institute of the California Institute of Technology to the Molecular Materials Research Center. **Funding:** This work was supported by a grant from the Human Frontier Science Program (RGP0014/2016), a France-Berkeley fund grant (FBF #2015-58) and an ANR grant (CLEARWING project, ANR-16-CE02-0012). **Author contributions:** Conceived and designed the experiments: AFP and NHP. Performed the experiments: AFP, RHS, EIC, KH. All authors contributed to analysis of the results, contributed reagents/materials/analysis tools and contributed to writing the paper. **Competing interests:** The authors declare they have no competing interests.

## Supplementary Materials

**Table S1. GC-MS relative proportions (mean ± standard deviation) of wing cuticular compounds isolated from *Greta oto***.

**Table S2. Spectrometry of *Greta oto* untreated and hexane treated clear wing regions and simulated reflectance spectra**.

**Movie S1. 3D projection of developing scales in a clear wing region 48 hours after pupal formation**. 3D projection and rotation of the same scales shown in Fig. 2F, 48 hours APF in a clear wing region. WGA (magenta) stains cell membranes and phalloidin (green) stains F-actin and DAPI (blue) stains nuclei. Short actin filaments have reorganized and formed smaller numbers of thick, regularly spaced parallel bundles just under the surface of the cell membrane. Scales alternate with future forked scales appearing as triangular shapes and longer future bristle-like shapes.

**Fig. S1.**
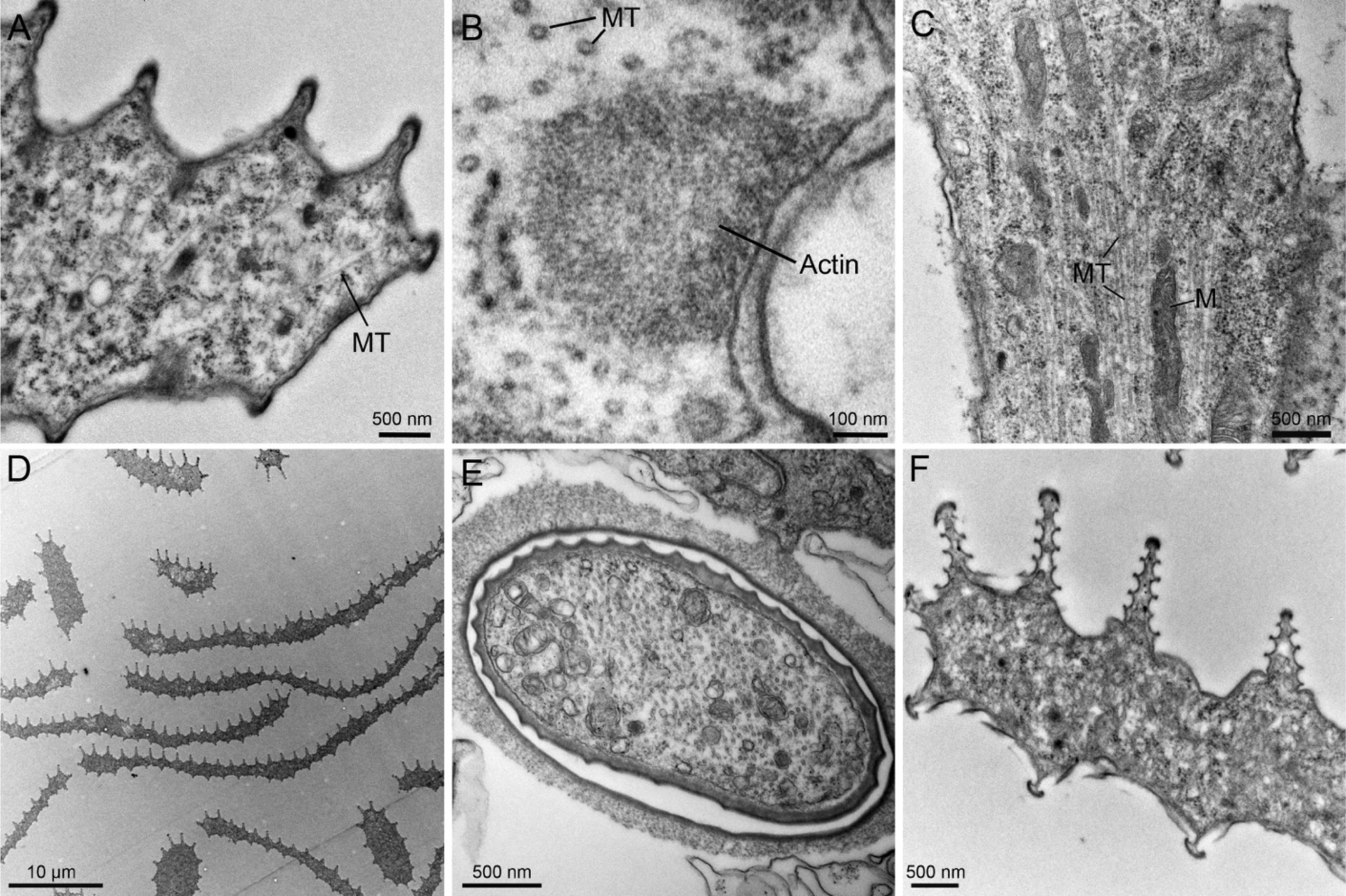
TEM micrographs of scales 72 hours (top) and 120 hours (bottom) after pupal formation. **(A)** TEM micrograph of a developing opaque scale 72 h APF, highlighting microtubule arrangement (MT). **(B)** Thick actin bundles contain dense, hexagonally packed F-actin filaments. **(C)** Basal region of a developing scale outgrowth and socket cell. Developing scales 72 h APF contain dense populations of microtubules (MT) and numerous internal organelles, including mitochondria (M), electron dense vesicles and free single ribosomes. **(D)** Transverse section of developing scales around 120 h APF, highlighting both flat and thin, bristle-like scale morphologies. Cross section near the **(E)** base and **(F)** distal region of scales 120 h APF, showing thickened cuticle and ridge morphologies.

**Fig. S2.**
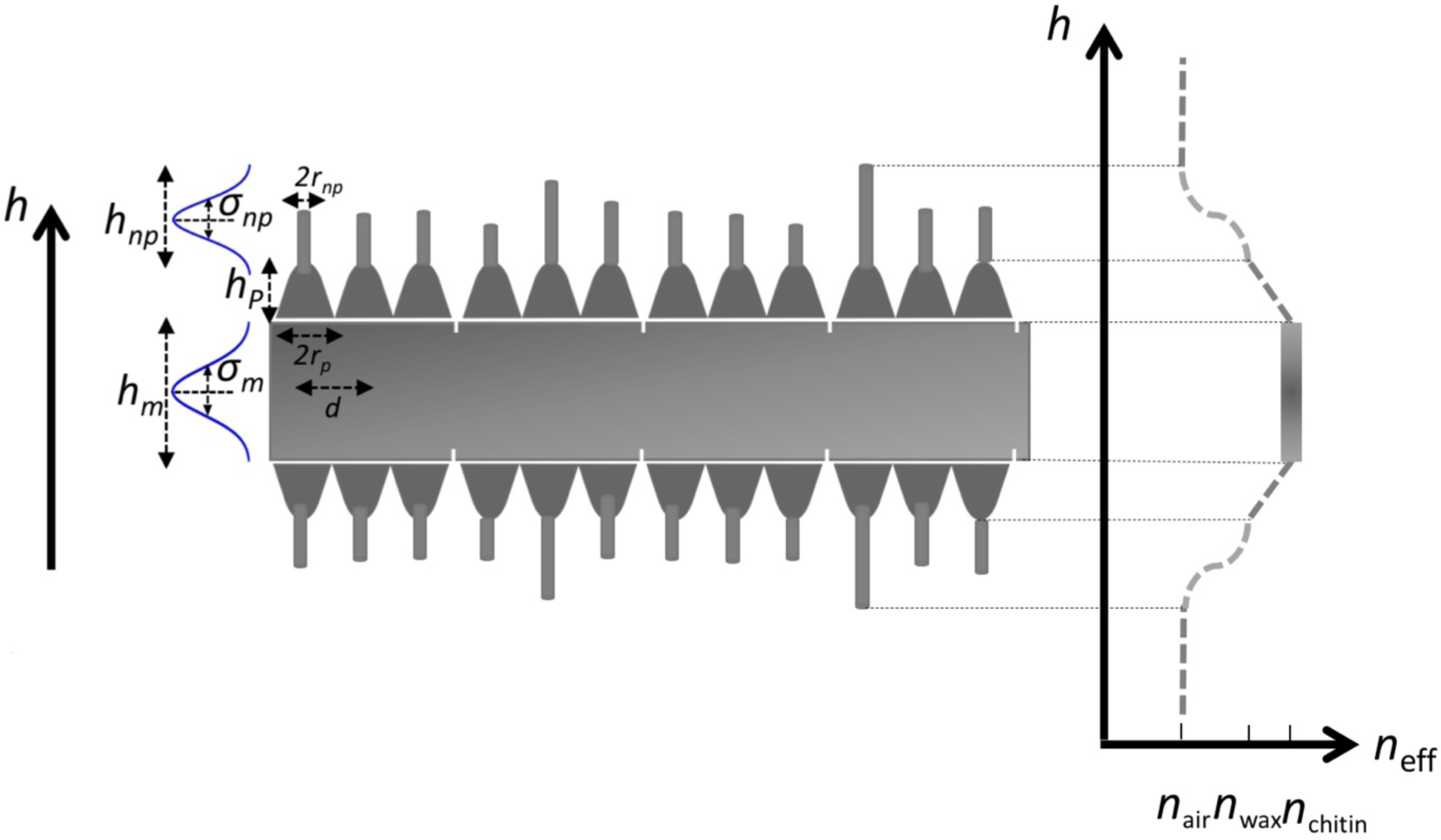
Optical modeling parameters and effective refractive index condition for untreated transparent wing of *Greta oto*. Schematic representation for the optical modeling parameters of wing membrane and surface nanostructures. Average distance between two nanostructures represented as *d*, conical shaped cuticular nipple nanostructures height as *h*_p_, wax-based irregular nanopillars radius as *r*_np_, mean height as *h*_np_ and variance *σ*_np_, and membrane thickness as *h*_m_ and variance *σ*_m_. Y-axis represents height *h* and X-axis represents effective refractive index condition of air (n_air_), chitin (n_chitin_), and wax (n_wax_).

**Fig. S3.**
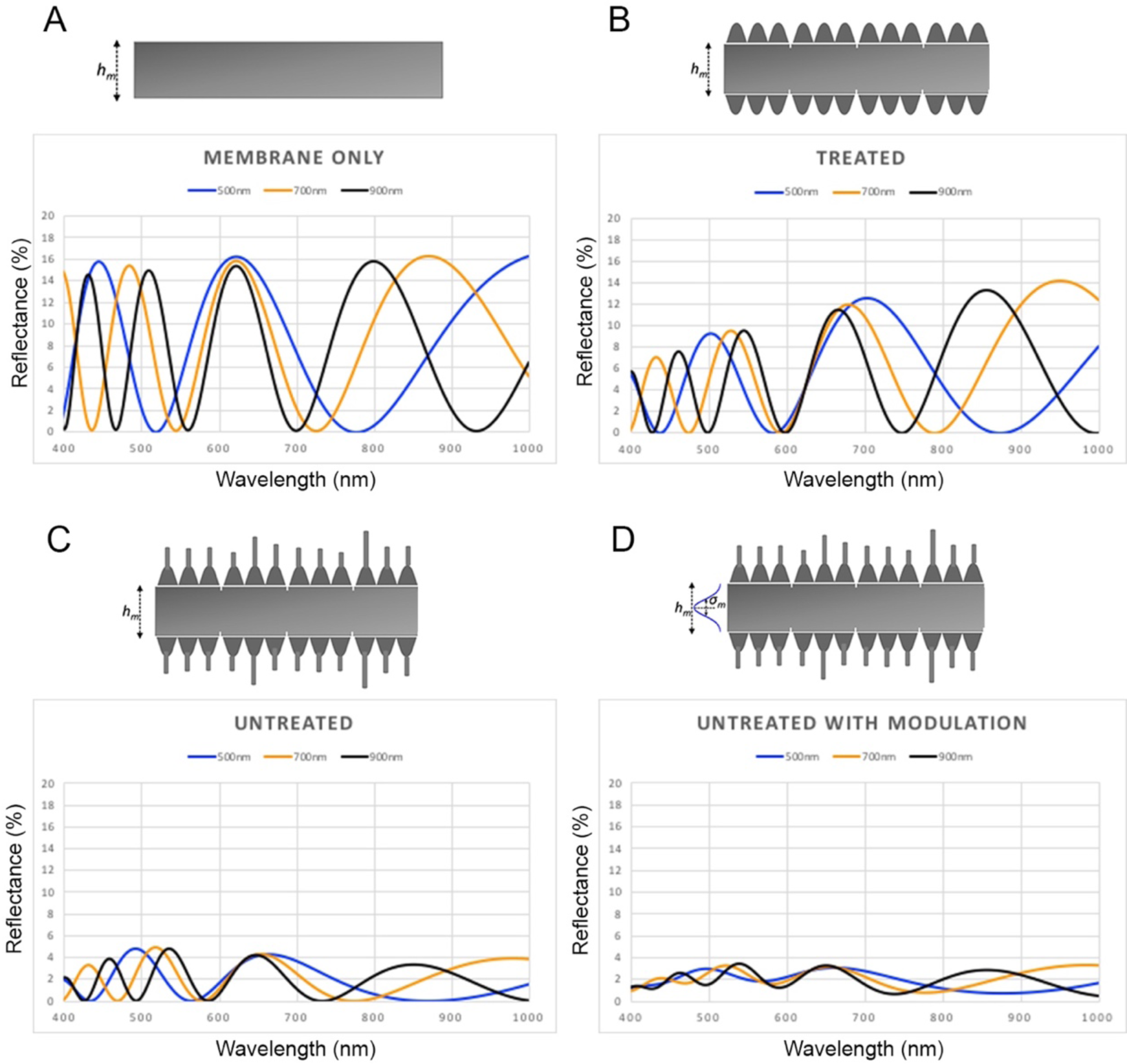
Optical simulations for mean membrane thickness and modulation of thickness under different wing architecture models. Simulation reflectance spectra of **(A)** Membrane only (lacking surface nanostructures) with varying mean membrane thickness. **(B)** Treated wings (containing cuticle-based nipple nanostructures but lacking wax-based irregular nanopillars) with varying mean membrane thickness. **(C)** Untreated wings (containing wax-based irregular nanopillars and nipple nanostructures) with varying mean membrane thickness and no modulation in thickness. **(D)** Untreated wings with variable mean membrane thickness and modulation of 43 nm variance in thickness.

